# Apical spectrin organizes cortical actin filament bundles to pattern *C. elegans* cuticle ridges

**DOI:** 10.64898/2026.03.05.709908

**Authors:** Prioty F. Sarwar, Trevor J. Barker, Ken Nguyen, Fung-Yi Chan, David H. Hall, Ana X. Carvalho, Meera V. Sundaram

## Abstract

The apical extracellular matrix can form elaborate three-dimensional structures on animal surfaces. To better understand the mechanisms that pattern and shape these structures, we focus on development of collagen-rich cuticle ridges (alae) in adult *C. elegans*. Previous studies suggested that longitudinal actin filament bundles (AFBs) in the lateral seam epidermis specify alae position through a mechanism that involves post-secretory matrix delamination. Here we identify additional components of this highly organized cortical actin network and show that loss of the apical βH-spectrin SMA-1 specifically disrupts organization of the two AFBs that would normally flank the site where the middle alae ridge forms. Correspondingly, *sma-1* loss, or mutation of its actin binding domains, also disrupts formation of the middle alae ridge. Ultrastructurally, *sma-1* mutants have expanded regions of matrix delamination that can explain middle ridge loss. Together, these data highlight the importance of apical spectrin for organizing a patterned actin network within epithelia and show that, via its effects on actin organization, spectrin can also change the extracellular matrix and its patterns on animal surfaces.

## Introduction

Animal surfaces are often decorated with three-dimensional apical extracellular matrix (aECM) structures such as pores, scales, denticles, or ridges [1–3]. Such aECM structures have characteristic spacing and morphology, and an important question is how they can be patterned and shaped within the extracellular environment.

Actin-dependent cues from the underlying epithelium can provide important spatial cues to initiate aECM patterning, but the mechanisms by which such cues are relayed to the matrix appear to vary. In some cases, actin filament bundles (AFBs) pattern aECM structures by altering the shape of underlying cells; for example, Drosophila denticle belts form when aECM is deposited on top of actomyosin-driven cellular protrusions that later recede, leaving the matrix behind [4,5]. Ridges in butterfly wing scales are also thought to form by actomyosin-driven membrane buckling [6]. In other cases, AFBs may determine sites of matrix secretion; for example, in budding yeast and Drosophila, actin rings or filaments position membrane-associated chitin synthase to pattern chitin-rich structures immediately above them [7–10]. In the case of the collagenous cuticle alae ridges of *C. elegans*, a third mechanism has been invoked that does not involve stable plasma membrane deformation or any apparent localized matrix secretion. Instead, ridges form in the intervals between parallel AFBs through a still poorly understood mechanism involving actomyosin-dependent matrix delamination [11].

*C. elegans* alae are longitudinal cuticle matrix ridges present over the lateral epidermis or “seam cells” of adults and other specific developmental stages (Fig. 1A) [1]. The three adult alae ridges form *de novo* as the new adult cuticle develops underneath the older non-ridged L4 larval cuticle (Fig. 1B). Although the mature alae are primarily collagenous in structure, proper formation of alae requires a different transient matrix, called the precuticle, present at their earliest stages of development [1,12]. Prior work showed that precuticle proteins, including the lipocalin LPR-3 and the Zona Pellucida (ZP) domain proteins LET-653 and NOAH-1, initially are secreted broadly over the apical surface of the seam syncytium, below the existing larval cuticle[11]. Subsequently, these precuticle proteins become patterned into bands corresponding to developing alae ridges or their intervening valleys (Fig. 1B,C), and they then help to organize the permanent matrix components that make up the mature adult alae.

**Figure 1.**
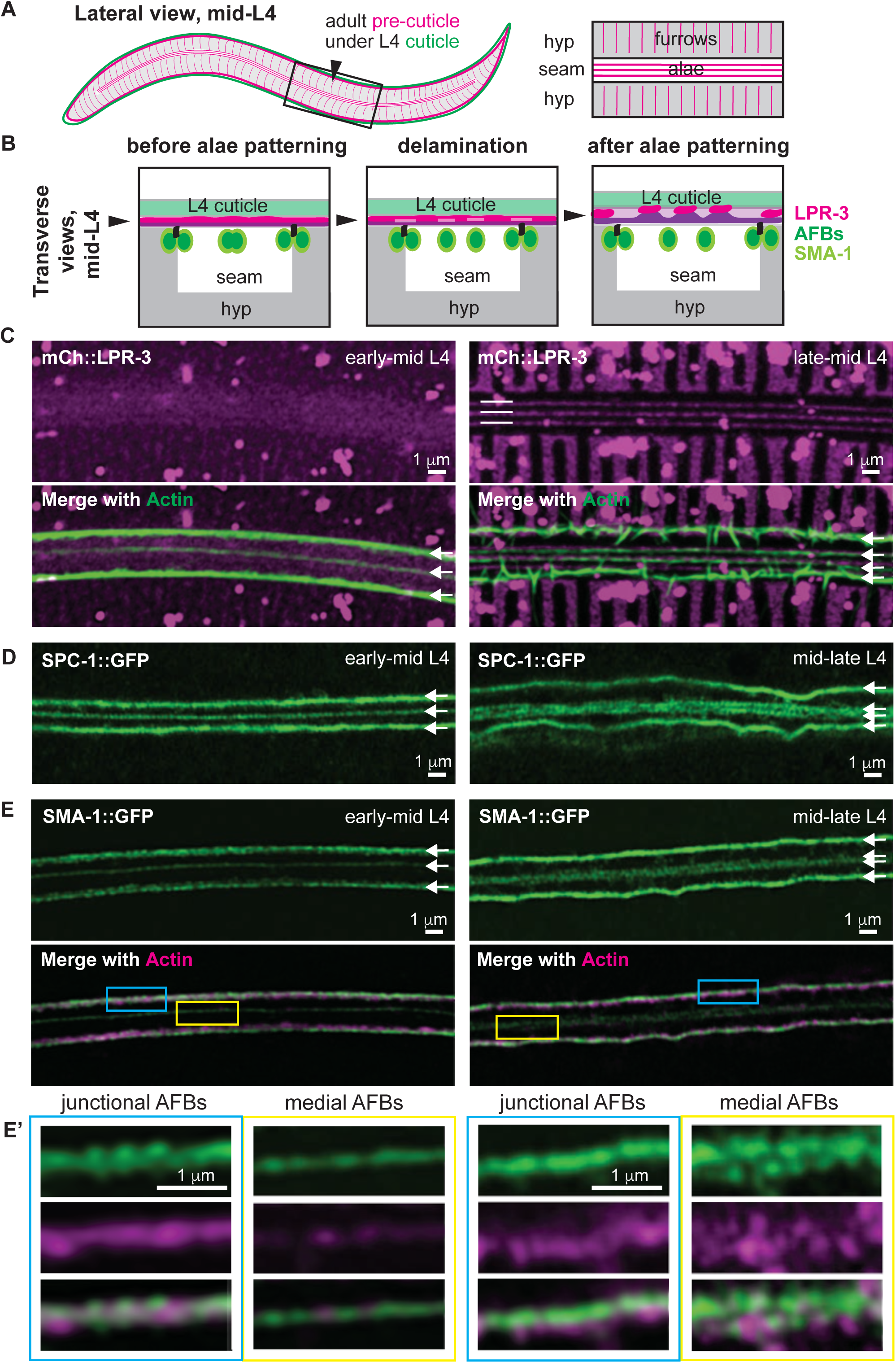
SMA-1/βH-spectrin associates with seam AFBs during alae patterning. (A) Lateral schematic of the adult precuticle (magenta) developing under the L4 cuticle (green).Black box indicates region shown at higher magnification in schematic at right and imaged in C-E. (B) Transverse schematics of the L4 seam syncytium, summarizing localization patterns of precuticle and cytoskeletal proteins during alae development. Alae form above the lateral seam syncytium, which sits within the larger hyp7 epidermal syncytium. (C-E) Super-resolution airyscan images of the apical seam at early (L4.3-L4.4, left) and late (L4.6-L4.7, right) mid-L4 substages, corresponding to before and after pre-cuticle alae patterning. The seam widens during alae patterning and the two medial AFBs become resolved [11]. (C) mCh::LPR-3 and seam AFBs (visualized using seam-expressed UTRNCH::GFP). Image at right is a Z-projection of six confocal slices; (D) SPC-1::GFP; (E) SMA-1::GFP and UTRNCH::DsRed. Boxed regions are shown at higher magnification in (E’). All images representative of at least n=10 specimens imaged.

The post-secretory patterning of the alae precuticle depends on the underlying actomyosin cytoskeleton [11]. Shortly after seam cell fusion to form left and right syncytia, and before precuticle patterning, four longitudinal AFBs form at the seam apical cortex and predict the sites of future alae valleys (Fig. 1B,C). Two of these seam AFBs are present at the junctions between the seam syncytium and the surrounding hyp7 epidermal syncytium, which has its own corresponding AFBs (Fig. 1B). The two remaining seam AFBs are present in the medial region of the seam, flanking the region where the middle alae ridge will form (Fig. 1B,C). RNAi knockdown of actin or non-muscle myosin (NM II) disrupted seam cell shape and precuticle patterning and ultimately led to disruptions in the mature alae. Ultrastructural data revealed that alae formation initiates as small horizonal slits or delaminations between layers of the precuticle aECM, without any accompanying stable deformation of the underlying seam plasma membrane (Fig. 1B). Like the seam AFBs, these matrix delaminations presaged where the valleys between alae ridges would form, while intervening regions of remaining matrix adhesion corresponded to the future alae ridges. These data suggested that longitudinal AFBs somehow relay information to the overlying matrix to trigger local delamination and the displacement of different matrix factors towards future ridge or valley regions. However, the very pleiotropic phenotypes seen after actin or NM II knockdown made it difficult to link specific changes in the cytoskeleton to changes in matrix structure.

Spectrins are a major component of the cortical membrane cytoskeleton, where they can help organize actin networks and link cortical actin to plasma membranes and specific transmembrane proteins [13]. Spectrins are also mechanosensitive and can help transduce internal cytoskeletal forces and/or external shear forces across the plasma membrane [14–16]. Therefore, they are excellent candidates for organizing the seam cytoskeletal network and/or relaying cytoskeletal cues to the apical membrane and developing matrix. Here we show that beta-heavy (βH) spectrin, along with the cytoskeletal cross-linker VAB-10/spectraplakin and intermediate filaments, are part of a highly organized but transient cytoskeletal network that assembles within the seam epidermis during the period of alae formation. SMA-1 is specifically required to organize the two medial-most AFBs within the seam and to pattern precuticle aECM factors to form the middle alae ridge. Ultrastructural data show expanded regions of matrix delamination in *sma-1* mutants at the earliest stages of alae patterning, demonstrating that SMA-1 and AFBs are required to focus delamination to specific regions. Finally, a tension-responsive LIM domain reporter reveals increased actin stress at remaining junctional AFBs, suggesting that altered seam mechanical properties may explain the altered balance between matrix delamination and adhesion.

## Results

### SMA-1/βH-spectrin and VAB-10/spectraplakin are part of a highly organized but transient cortical cytoskeletal network within the seam epidermis

To identify molecular players that pattern alae development, we first screened endogenous fluorescent fusions to known actin-binding proteins and other cytoskeletal factors, looking for AFB-related localization patterns (Table S1). This screen identified SMA-1/βH-spectrin and VAB-10/spectraplakin, amongst others, as part of the apical cytoskeletal network within the seam epidermis (Fig. 1 and Fig. 2).

**Figure 2:**
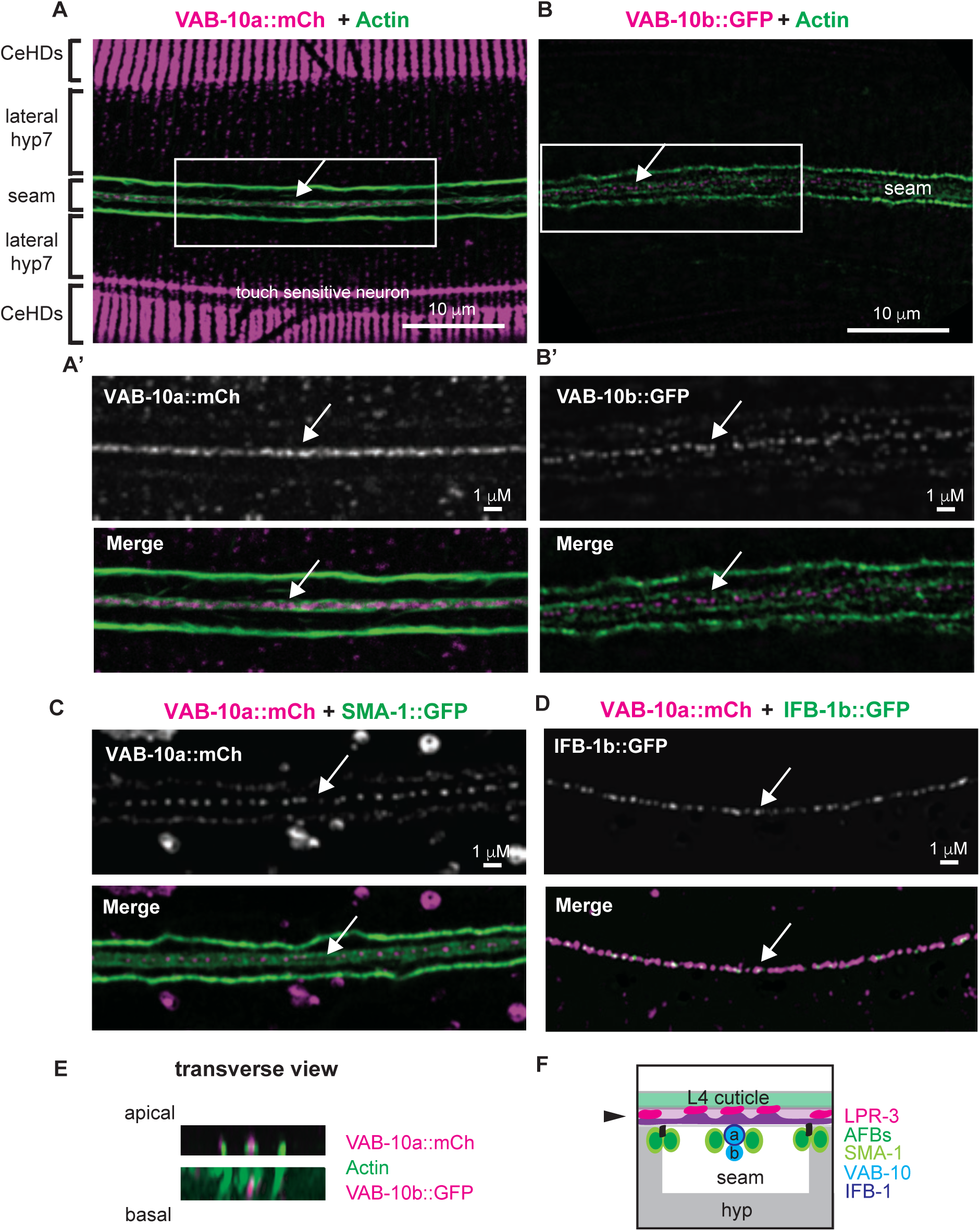
VAB-10/spectraplakin and IFB-1 localize between seam medial AFBs. (A-D) Super-resolution airyscan images of the apical seam at the L4.5-L4.7 substages. (A, A’, C) VAB-10a::mCh and (B,B’) VAB-10b::GFP form a line of puncta between the two medial seam AFBs. VAB-10a prominently marks CeHDs, as previously described [21], but VAB-10b was not detected at CeHDs at this stage. VAB-10a is also present in faint circumferential bands of puncta in the lateral portion of hyp7, similar to SMA-1 (Fig. S1) and actin [11]. (D) VAB-10a::mCh and IFB-1b::GFP puncta are found interspersed together at the medial seam. (E) Orthogonal (transverse) views of the animals in A and B. VAB-10a appears slightly apical to AFBs while VAB-10b appears slightly basal. (F) Transverse schematic of the L4 seam syncytium. VAB-10 and IFB-1b are found below the developing middle alae ridge. All images representative of at least n=10 specimens imaged.

Spectrins usually form heterotetramers with two alpha and two beta subunits [13]. *C. elegans* has one alpha spectrin (SPC-1), one beta heavy chain spectrin (SMA-1), and one canonical beta spectrin (UNC-70) (see Fig. 3A). Consistent with prior studies in other tissues [17–19], we observed SMA-1::GFP apically within the seam, whereas UNC-70::GFP was basolateral; SPC-1::GFP was present in both domains, consistent with SPC-1 functioning with both beta subunits (Fig. 1D,E and Fig. S1). The known spectrin binding protein, UNC-44/ankyrin, resembled UNC-70 in its basolateral localization (Fig. S1). Both the SMA-1 and SPC-1 apical patterns resembled the actin pattern, with three longitudinal bands in the L4.3-L4.4 substages (“early mid-L4”, when the seam is at its narrowest) that resolved to four bands in the L4.5-L4.7 substages (“late mid-L4”, when the seam widens and developing alae first appear) (Fig. 1D,E). SMA-1 and actin co-labelling showed that they are closely associated, though not precisely colocalized (Fig. 1E). Similar to actin, the SMA-1/βH-spectrin and SPC-1/α-spectrin bands were present only transiently during the mid-L4 stages (L4.3-L4.7, Fig. 1D,E) and dispersed before the molt to adulthood (Fig. S1). Thus, these cytoskeletal structures correlate well with the time when alae ridges are being patterned and built.

**Figure 3:**
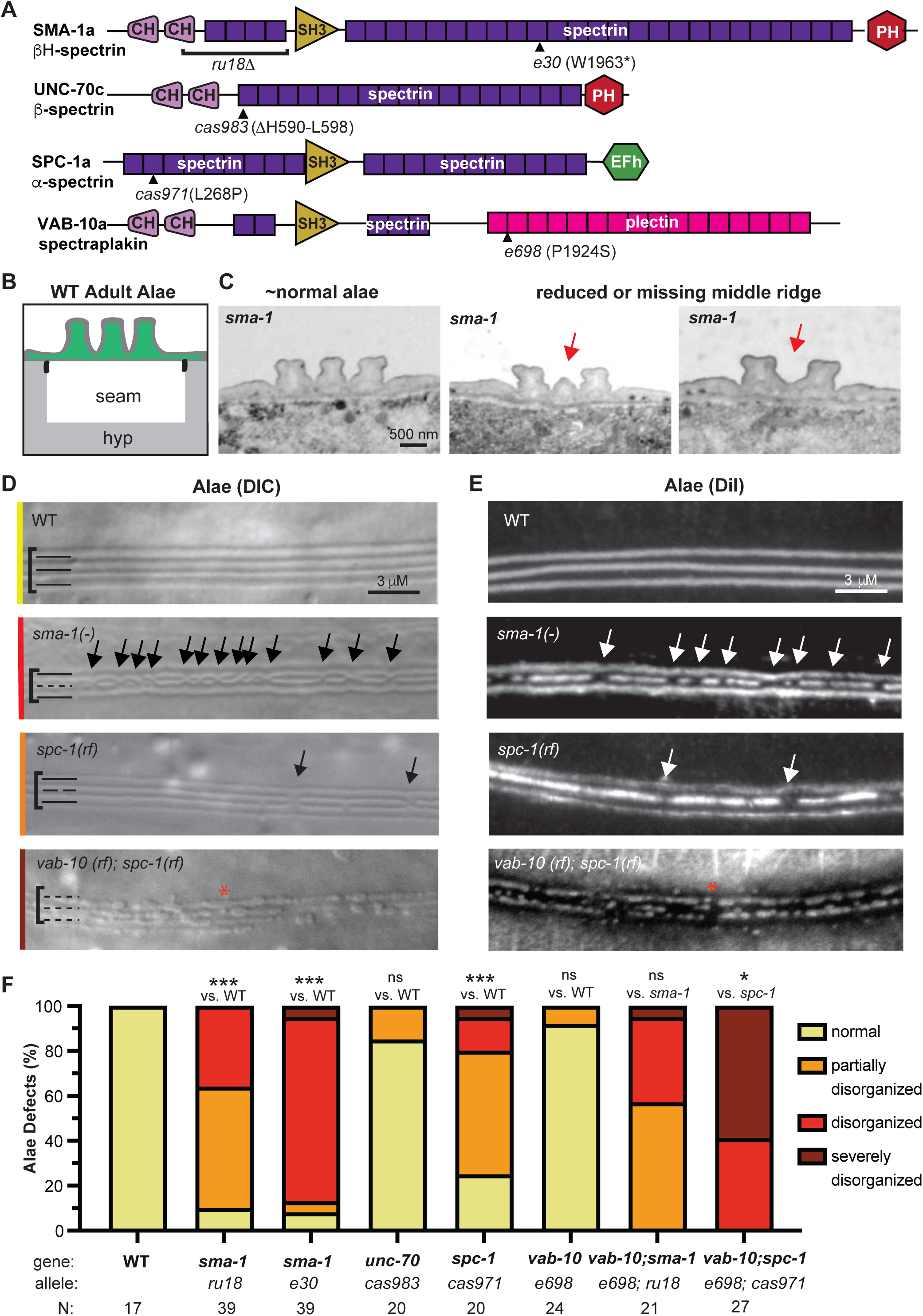
Apical spectrin mutants have defects in the middle alae ridge. (A) Protein schematics highlighting functional domains of the *C. elegans* spectrin and spectraplakin proteins and mutant lesions. The specific isoforms drawn are indicated. (B) Transverse schematic of the seam and alae in wild-type (WT) adults. (C) Archival TEM images of *sma-1(e30)* adults (a kind gift from M. Buechner). Transverse sections from different body regions show some areas with normal alae (left) and other areas with an under-developed (middle) or absent (right) middle ridge (indicated with red arrows). (D) Representative images of the young adult alae of indicated strains, as viewed by Differential Interference Contrast (DIC) microscopy. Brackets encompass the alae region. Solid lines at left indicate intact alae ridges while dashed lines indicate disrupted alae. Colored lines on the left correspond to how the image was categorized in (F). (E) Young adult alae viewed by epifluorescence after staining with the lipophilic dye DiI. Arrows point to prominent middle alae breaks. Red asterisks indicate when all three alae ridges show noticeable defects. (F) Phenotypic penetrance and expressivity of alae defects in single or double mutants. The *vab-10a(e698)* and *spc-1(cas971)* missense alleles showed a strong genetic interaction. *** p<0.0001, Fisher’s Exact Test (normal vs. abnormal) when compared to WT and ** p<0.001 (severe vs. other) when compared to *spc-1* single mutant. ns, not significant.

VAB-10/spectraplakin is a cytoskeletal linker thought to connect actin to intermediate filaments (IFs) and microtubules (MTs) [20]. It has been studied mainly for its role in *C. elegans* hemidesmosomes (CeHDs), which span narrow regions of the hyp7 epidermis to link the cuticle to underlying body muscle [21]. Using endogenous fusions (Table S1), we also observed VAB-10 in other regions of hyp7 and in the seam (Fig. 2A). Within the seam, the two major splice isoforms VAB-10a (IF binding) and VAB-10b (MT binding) each formed a prominent line of medial puncta running between the two medial AFBs and SMA-1/βH-spectrin (Fig 2A-C). As previously reported in embryonic CeHDs [21,22], VAB-10a appeared closely associated with the intermediate filament IFB-1b (Fig. 2D) and it localized slightly more apically than VAB-10b when compared to AFBs (Fig. 2E,F). Thus, VAB-10 and IFs run beneath the region where the middle alae ridge will form.

Several other CeHD components (VAB-19, LET-805) were not detected in the medial region of the mid-L4 seam (Table S1), suggesting the lack of a canonical CeHD structure at this site and consistent with the absence of muscle under most of the seam epidermis. The apical CeHD component MUP-4 (a matrillin-related transmembrane protein [23]) was found near VAB-10 in a single band in the midbody, but it appeared at a later stage than VAB-10 and, unlike VAB-10, it did not extend beyond the midbody (Fig. S2). Furthermore, MUP-4 appeared more basal than VAB-10a, suggesting it might be present at hyp7 and/or vulva cell membranes below the seam, near where CeHDs will later form above vulva muscle attachments (Fig. S2). We conclude that, at least outside the midbody, the seam apical VAB-10 structure does not correspond to a classical CeHD but instead represents a distinct cortical arrangement of VAB-10 and IFs.

### *sma-1/*βH-spectrin mutants have defects in the middle alae ridge

To test for functional roles of apical spectrin, we examined mutants for two different *sma-1* alleles, including the frameshift and protein null *ru18* and the nonsense allele *e30* (Fig. 3A), both of which are viable [17]. In *wild-type* (*WT*) young adults, the three alae ridges each have a roughly rectangular shape when viewed in transverse sections by transmission electron microscopy (TEM) (Fig. 3B,C). Alae can also be visualized as longitudinal bands using differential interference contrast (DIC) microscopy or by staining with the lipophilic dye DiI (Fig. 3D,E). The *sma-1* alleles caused defects specifically in the middle alae ridge, which appeared missing or under-developed in some regions, while the two outer alae ridges appeared largely normal (Fig. 3C-E).

Using DIC imaging, we categorized mutant severity based on whether the middle ridge was consistently disrupted across the mid-body (“disorganized”) or only occasionally disrupted with long intervening stretches of normality (“partially disorganized”). Rarely, both middle and outer ridges were disrupted (“severe”). Surprisingly, the two *sma-1* alleles showed differing severity and penetrance, with *ru18* being less severe than *e30* (Fig. 3D-F). It is possible that *sma-1(e30)* produces some truncated protein, causing a greater steric disruption to the spectrin network. However, we can’t exclude the possibility of a linked modifier. All further experiments were done with the null allele *sma-1(ru18)*.

*spc-1* and *unc-70* null mutants are early larval lethal [18], so we examined only partial reduction of function (rf) missense mutants (Fig. 3D-F) [24]. *spc-1(cas971rf)* caused middle alae defects similar to, but slightly less severe than, those caused by loss of *sma-1*, while most *unc-70(cas983rf)* mutants appeared normal (Fig. 3D-F). We conclude that the apical spectrin network (SMA-1 + SPC-1) is important for patterning the middle alae ridge.

### A *vab-10a*/spectraplakin mutant genetically interacts with *spc-1*/α-spectrin

*vab-10* null mutants are embryonic lethal [21], so to test for functional roles of spectraplakin we examined RNAi-treated animals and multiple partial reduction of function mutants; however, none showed any alae defects (Table S2). *Ifb-1b* mutants were also mostly normal (Table S2). We then asked if *vab-10* showed any genetic interactions with spectrins. The *vab-10a*-specific missense allele *e698* (Fig. 3C) did not enhance the defects of *sma-1(ru18)* mutants, but it did strongly enhance *spc-1(cas971rf),* causing severe defects across all three alae ridges (Fig. 3D-F). We conclude that VAB-10 cooperates with spectrins to pattern all three alae ridges, and therefore, like actin, VAB-10 and α-spectrin play broader roles than SMA-1.However, since VAB-10 appears to play an accessory role, or one that is masked by its pleiotropy, we focused further studies on SMA-1/βH-spectrin.

### SMA-1/βH-spectrin functions within the seam syncytium to affect alae patterning

Since both the seam epidermis and hyp7 have been implicated in alae patterning [11], and SMA-1 is expressed in both tissues (Fig. S1) [25], we used tissue-specific RNAi to test if *sma-1* acts in the seam to affect alae patterning (Fig. 4). These experiments used an RNAi-deficient *rde-1* mutant strain with transgenic rescue of *rde-1* under the seam-specific promoter *elt-5pro* [11]. As a control, we also included a strain with transgenic rescue of *rde-1* under the Pnp specific promoter *lin-31pro* [26], since both Pnp-derived cells and seam-derived cells ultimately fuse with hyp7 [27] and might therefore confer some RNAi sensitivity there. Seam-specific RNAi knockdown of *sma-1* caused alae defects similar to those seen in whole-animal RNAi positive controls, while Pnp-specific RNAi knockdown showed very few defects, similar to *rde-1* negative controls (Fig. 4A,B). We conclude that *sma-1* functions within the seam syncytium to pattern the middle alae ridge.

**Figure 4.**
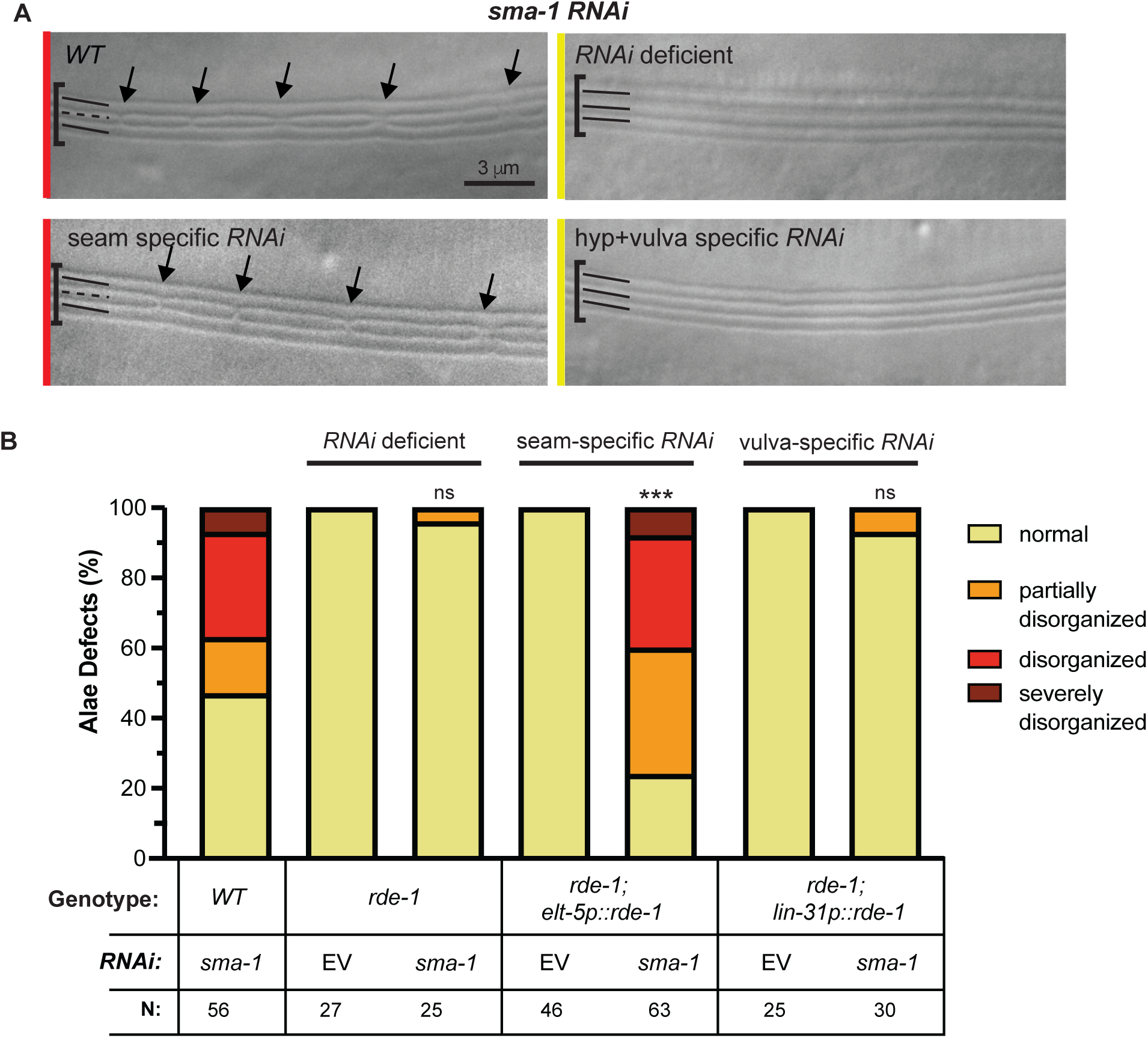
*sma-1* acts within the seam to pattern the middle alae ridge. (A) Representative DIC micrographs of the adult alae after *sma-1 RNAi* treatment in: i) *WT* (N2) ii) *rde-1* (*RNAi* deficient) iii) *rde-1*; *elt5pro*::*rde-1* (seam specific *RNAi*) and iv) *rde-1*; *lin-31pro*::*rde-1* (hyp7+vulva specific RNAi). Colored lines on the left correspond to how the image was categorized in (B). Arrows are pointing to significant breaks in the alae. (B) Quantification of alae defects in *sma-1* RNAi treatment condition versus empty vector (EV) control. ***p<0.0001, ns (not significant) using Fisher’s Exact Test (normal vs. abnormal) when compared to EV.

### *sma-1* actin binding domains (ABDs) are important for normal alae patterning

To better understand the mechanism by which SMA-1 acts, we took advantage of available domain-specific *sma-1* mutants [28] (Fig. 5A). Deletions removing the SMA-1 pleckstrin homology (PH) or SH3 domains did not cause any alae defects (Fig. 5B,C), suggesting that PH-mediated interactions with the plasma membrane and SH3-mediated signaling are not critical to SMA-1 function. A deletion removing 11 of the 29 spectrin repeats caused some defects, but these were relatively modest (Fig. 5B,C), suggesting a limited role for these repeats. On the other hand, two deletions removing parts of the actin binding calponin-homology (CH) domains caused middle ridge defects as severe as those in *sma-1(e30)* mutants and more severe than those seen in *sma-1(rh18)* null mutants (Fig. 5B,C; compare to Fig. 3F). Importantly, these latter alleles both reduce SMA-1 protein levels to varying degrees but do not eliminate expression, as assessed by Western blot, with ΔABD2 retaining more expression thanΔABD1 [28]. Nevertheless, the phenotypes of these two alleles were indistinguishable (Fig. 5C). These results suggest that actin binding is critical for SMA-1-dependent alae patterning.

**Figure 5.**
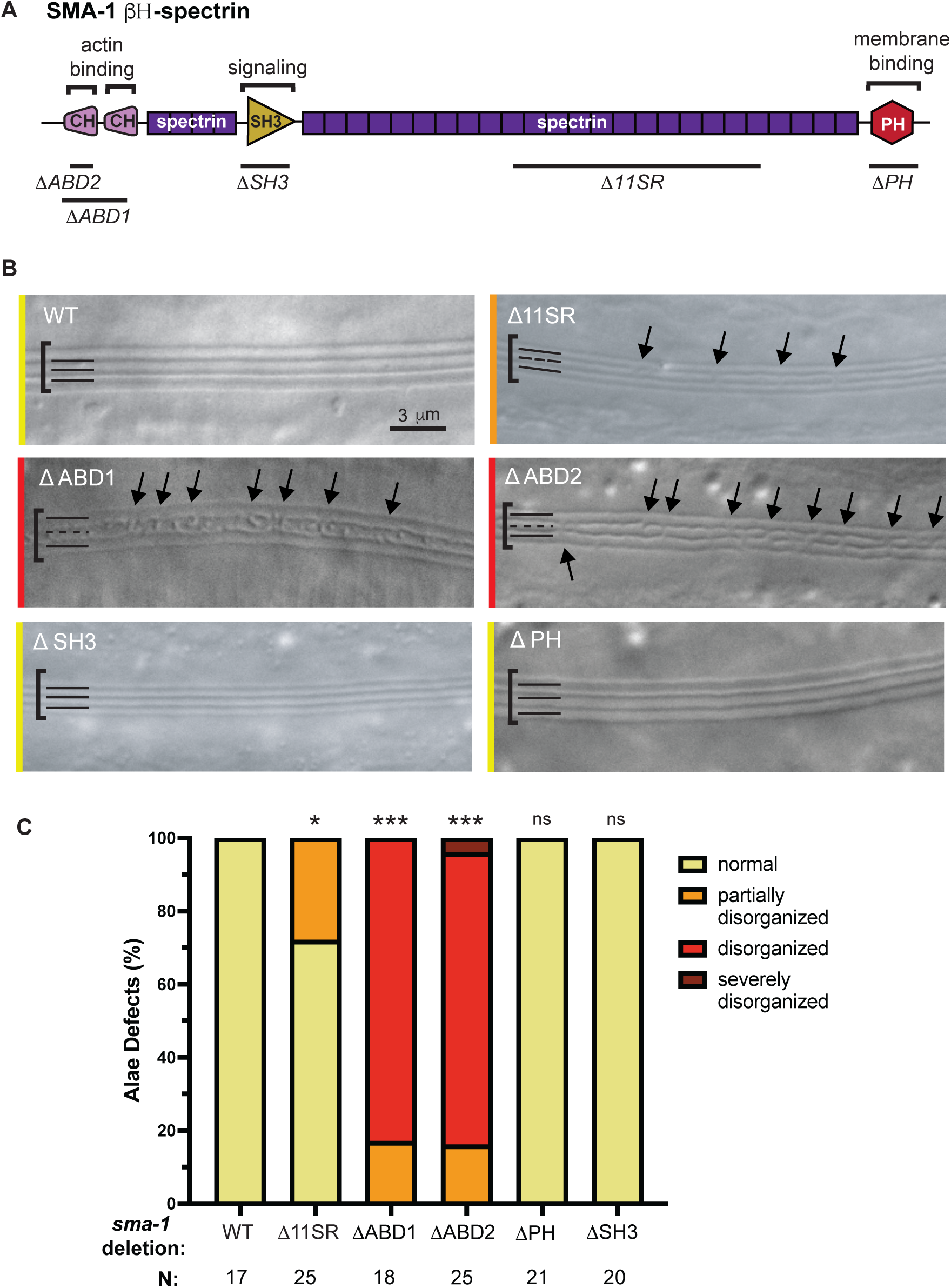
The SMA-1 actin binding domain (ABD) is required to pattern the middle alae ridge. (A) SMA-1 protein schematic highlighting functional domains and domain-specific deletions. (B) Representative DIC images of the alae phenotype in five *sma-1* domain mutants. Colored lines on the left correspond to how the image was categorized in (C). Black arrows indicate prominent alae breaks. Brackets encompass the alae region. Solid line indicates an overall linear alae appearance while dashed lines indicate disrupted alae. (C) Quantification of alae defects. *p<0.01, ***p<0.0001, ns (not significant) using Fisher’s Exact Test (normal vs. abnormal) when compared to WT.

### *sma-1* loss specifically disrupts organization of the medial seam AFBs

To ask at what step SMA-1 acts to pattern alae, we examined molecular markers for the precuticle, apical junctions, and the cytoskeleton. Consistent with their observed alae phenotype (Fig. 3), *sma-1* mutants showed mis-patterning of the pre-cuticle matrix factor LPR-3, specifically disrupting the developing middle alae ridge (Fig. 6A). Therefore, *sma-1* acts at a step before pre-cuticle patterning. VAB-10a medial puncta were still present, though slightly more disorganized, indicating that SMA-1 is dispensable for VAB-10 recruitment (Fig. 6B).Furthermore, unlike we previously reported for actin or NM II knockdown [11], *sma-1* loss did not disrupt apical junctions or the continuity and overall shape of the seam across the L4 stage (Fig. 6C,D and Fig. S3). SPC-1 still localized to the hyp7-seam border, likely due to its interactions with UNC-70/β-spectrin, but *sma-1* mutants completely lacked apical SPC-1 at the medial seam (Fig. 6C). Finally, *sma-1* mutants still had normal junctional AFBs and NMY-2/NM II, but medial AFBs were much reduced, with apical actin still present in some longitudinal structures but appearing much fainter and less coherent (Fig. 6D-F). Together with the requirement for the SMA-1 actin binding domains (Fig. 5), these data indicate that SMA-1/βH-spectrin promotes proper medial AFB organization and/or stability to pattern the pre-cuticle at the middle alae ridge.

**Figure 6.**
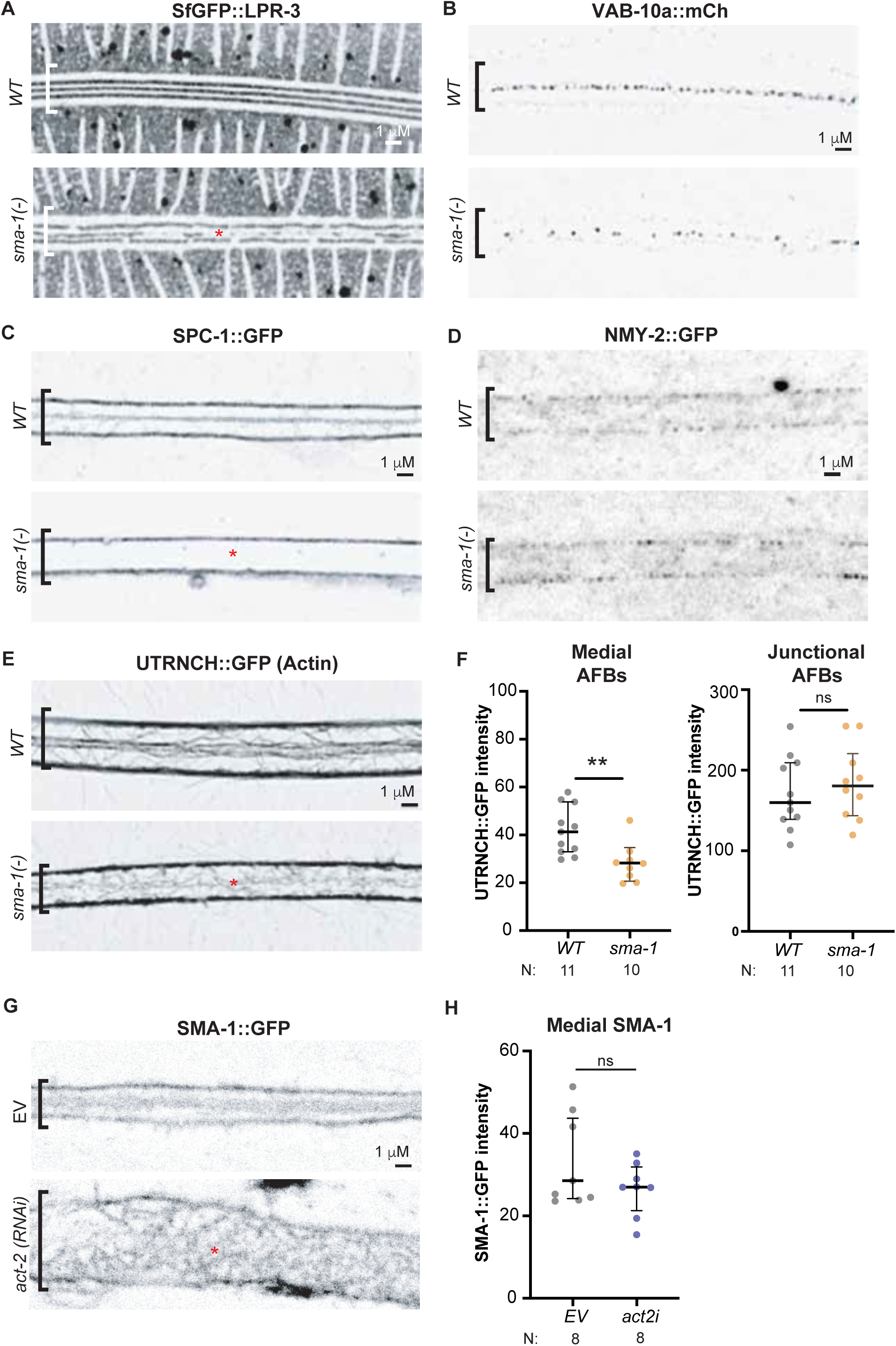
*sma-1* mutants have reduced medial AFBs. (A-E) Comparison of fluorescently tagged protein localization in a WT versus *sma-1(-)* background. All images are Airyscan processed, shown in inverted greyscale, and are representative of at least n=10 specimens imaged. Red asterisks indicate abnormalities. (A) SfGFP::LPR-3 precuticle marker reveals fragmentation of the developing alae ridge in *sma-1*(-) mutants. (B) VAB-10a::GFP puncta still mark the medial seam. (C) SPC-1::GFP is lost from the apical region in *sma-1(-)* mutants. (D) NMY-2::GFP is mainly detected at junctional AFBs in both *WT* and *sma-1* mutants. The signal at medial AFBs is too low for reliable assessment. (E) Seam AFBs visualized by *elt-5pro::*UTRNCH::GFP are reduced and less coherent in the medial region of *sma-1*(-) mutants, while junctional AFBs remain intact. (F) Quantifications of UTRNCH::GFP intensity from (E).Each dot represents the average intensity from three measured regions, spaced 5 µm and 10um apart, within a single animal. **p<0.001, Mann Whitney U test. ns, not significant. (G) Confocal images of SMA-1::GFP localization in empty vector (EV) control versus *act-2* RNAi background shows irregular and tortuous patterning at both the hyp7-seam junctions and middle of the seam. (H) Quantification of the medial apical SMA-1::GFP signal from (G). Despite the effect on patterning, no significant difference (ns) in the amount of fluorescent signal was found by Mann-Whitney U test.

As expected, *act-2*/actin RNAi greatly disrupted SMA-1 patterning at both the junctional and medial regions, though substantial amounts of SMA-1 remained near the apical cortex, likely due to residual actin (Fig. 6G,H). Therefore, SMA-1 and actin are mutually dependent for setting up or stabiilizing the organized medial bands that ultimately pattern the middle alae ridge.

### Ultrastructure reveals expanded regions of matrix delamination in *sma-1* mutants

We predicted that the reduction in medial AFBs might reduce matrix delamination in *sma-1* mutants. On the contrary, transmission electron microscopy showed enlarged regions of delamination (Fig. 7A-D). While medial delaminations were on average <200 nm wide in wild type, they were >300 nm wide in *sma-1* mutants (Fig. 7D). In some cases, the two medial delaminations were very close together or merged into one, without an intervening region of matrix adhesion, explaining loss of the middle alae ridge (Fig. 7B,C). In older L4 specimens, we also observed cases where three alae ridges had begun to form, but the middle ridge was underdeveloped and appeared to have prematurely lost adhesion to the old (L4) cuticle (Fig.7E). This latter phenotype corresponds well to the underdeveloped middle ridge also observed in TEM of *sma-1* adults (Fig. 1C). At all stages, the seam apical membrane remained flat in *sma-1* mutants. Apical vesicles were frequently observed, but these did not correlate with slit position and there was no apparent difference in their abundance between genotypes. These results indicate that SMA-1 and medial AFBs are not required for matrix delamination *per se*, but rather influence the spatial distribution of delamination vs. adhesion.

**Figure 7.**
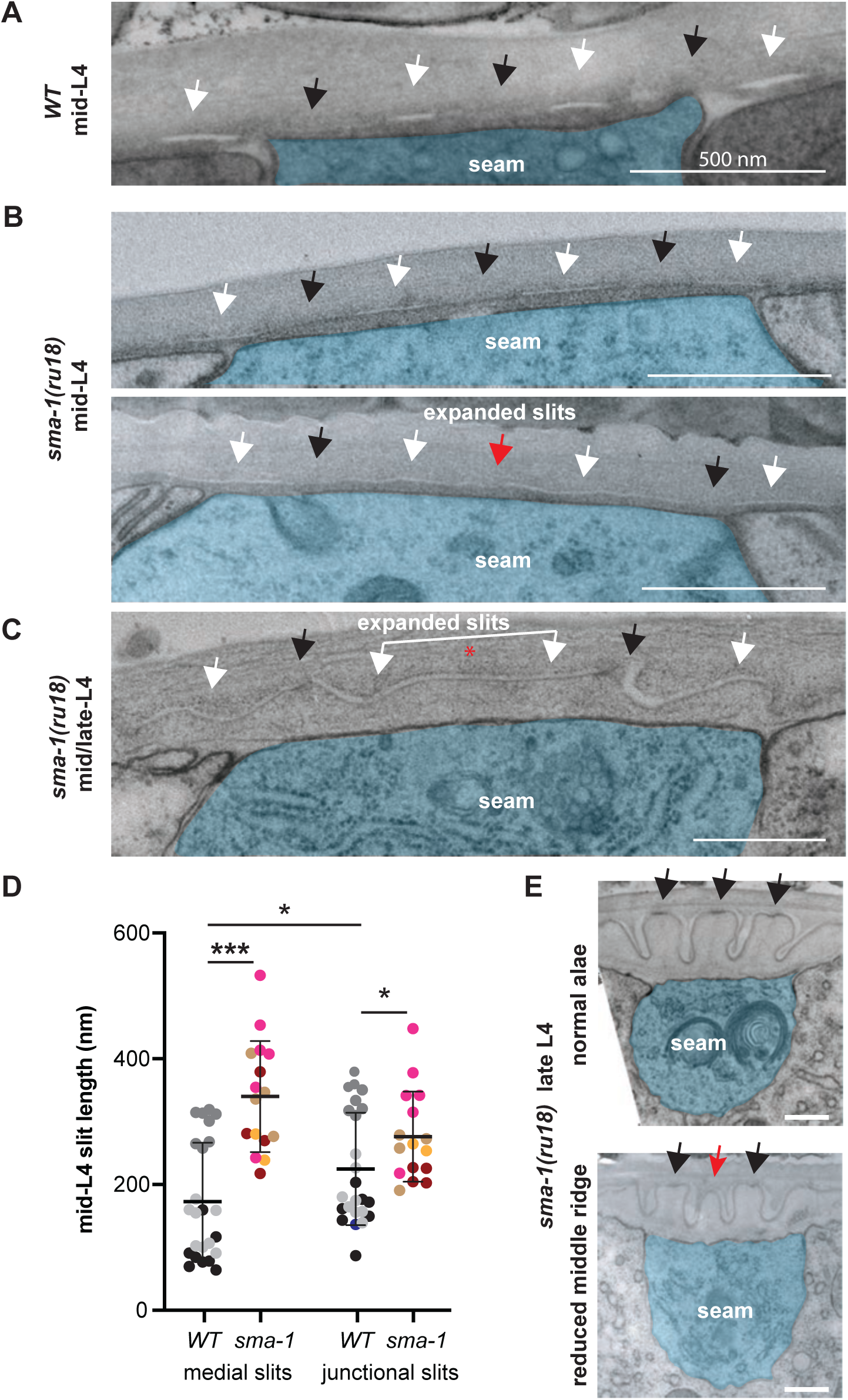
*sma-1* mutants have expanded regions of matrix delamination. (A-C) TEM micrographs of transversely-sectioned mid-L4 larvae. Seam cell is false-colored in blue. White arrows indicate matrix delaminations or slits, and black arrows indicate regions of remaining matrix adhesion. Red arrow and asterisk indicate regions where adhesion has been nearly or fully lost, respectively. Scale bars: 500 nm. (A) wild-type (N2 1455-F). (B,C) *sma-1(ru18).* In B, the top image (AZ30-1B) appears similar to WT, while the bottom image (AZ30-6) shows expanded slits. The animal in C (AZ30-7) is slightly older and its outer alae ridges have begun to form and contain electron dense material at their tips (black arrows); however no middle ridge or density is present. (D) Quantification of slit length in mid-L4 larvae. Each dot represents a single slit measurement (with 4 measurements per image) and colors indicate the different specimens (n=3 WT and n=4 *sma-1(ru18)*). Where possible, measurements were made of slits on both the left and right sides of two separate body regions. Light gray, N2 1455-F; dark gray, N2 1455-B; black, N2 1455-H; maroon, AZ30-1A; light brown, AZ30-1B; gold, AZ30-5; magenta, AZ30-6B). Note that the *sma-1* specimen in C was excluded from this analysis because of its slightly older age, but it shows a similar trend. *p<0.01, ***p<0.0001, Mann Whitney U test. (E) Two regions of the same *sma-1(ru18)* late L4 larva (AZ30-3). At the top, all three alae ridges have formed normally (black arrows), while at the bottom the middle ridge is smaller and lacks electron dense material at its tip (red arrow).

### *sma-1* mutants have altered seam mechanics

Since *sma-1* mutants have reduced medial AFBs, we hypothesized that the seam cortical network would experience altered forces. To test this, we used a previously described tension reporter that is based on the ability of LIM domain proteins to bind stressed actin [29,30]. TES-1/testin is a LIM domain protein expressed in the seam but not the surrounding hypodermis, and its interaction with actin is facilitated by removal of an auto-inhibitory PET domain (Fig. 8A). In wild type, mNG::TES-1(ΔPET) was consistently present at seam junctional AFBs and (to a lesser degree) at medial AFBs throughout L4, suggesting fairly constant actin stress levels within this tissue (Fig. 8B,C). Strikingly, in *sma-1* mutants, mNG::TES-1(ΔPET) accumulated twice as highly at junctional AFBs (Fig. 8B,C,C’). Nevertheless, we did not see a corresponding increase in NMY-2::GFP, which was faint along junctional AFBs and nearly undetectable at medial AFBs in both WT and *sma-1* animals (Fig. 6D). Junctional actin levels also were similar between WT and *sma-1* animals (Fig. 6E,F), so the increase in TES-1 signal suggests an increase in actin stress at that location. These experiments suggest that medial AFBs help distribute forces within the seam syncytium and show that *sma-1* mutants have altered force distributions that could contribute to the altered patterns of matrix delamination.

**Figure 8.**
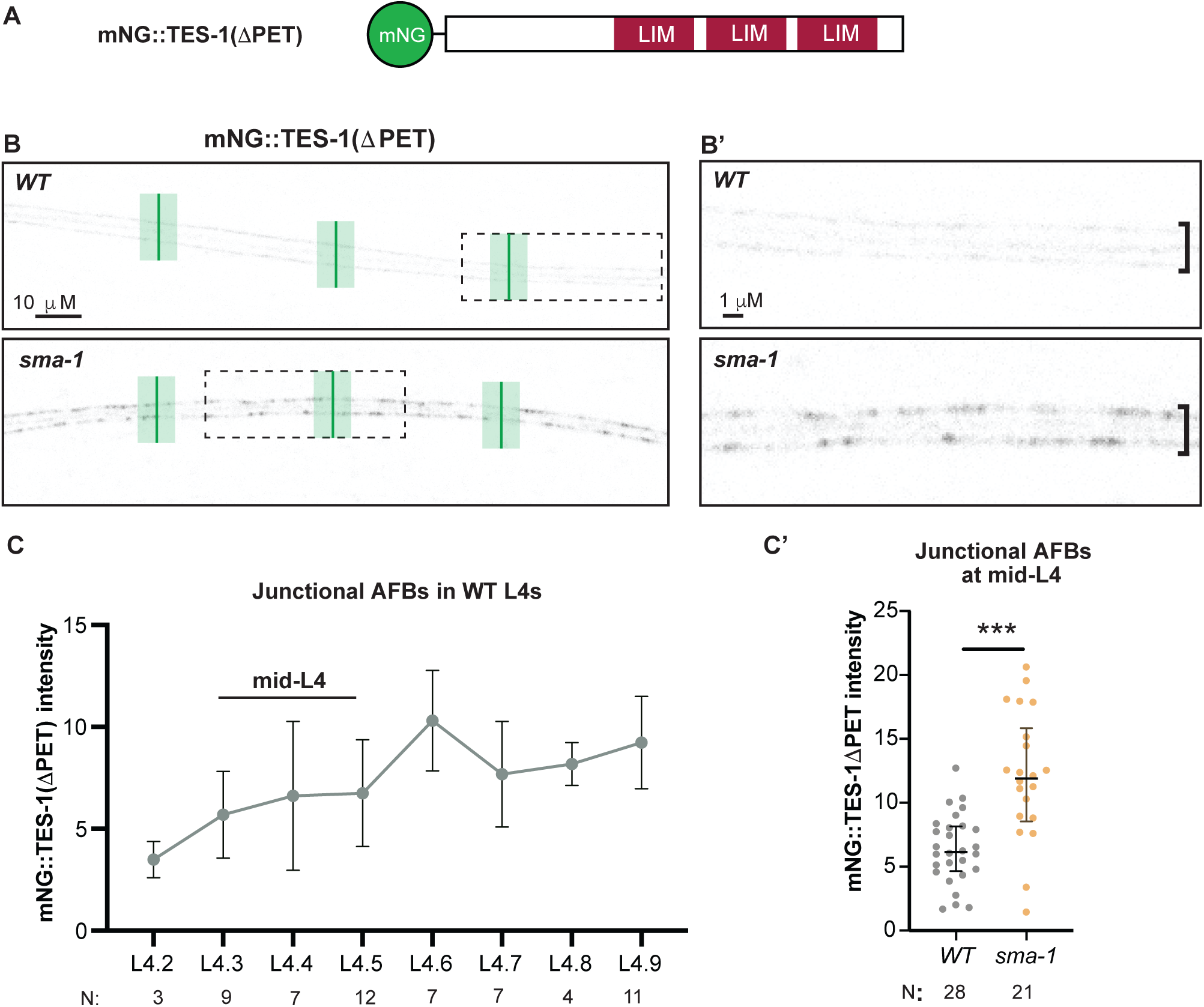
*sma-1* mutants have altered seam mechanical stress. (A) Schematic of mNeonGreen (mNG) tagged TES-1ΔPET, previously used as an AFB tension reporter [29]. (B,B’) Representative single confocal slices of mNG::TES-1ΔPET in WT vs *sma-1(-)* background at mid-L4 (L4.3-L4.5), shown in inverted greyscale. In (B), green bars indicate the three locations per image where peak junctional intensity was measured. Boxed regions are magnified in B’. In *sma-1(-)* mutants, mNG::TES-1ΔPET signal was consistently elevated in a patchy pattern along the length of the seam. (C, C’) Quantification of mNG::TES-1ΔPET intensity as shown in (B).Each dot represents the average from all measurements within a single animal. **p<0.001,***p<0.0001, Mann Whitney U test.

## Discussion

Spectrins organize and stabilize diverse cortical actin arrangements to shape and protect plasma membrane domains in red blood cells, neurons, and epithelia [13,16]. Here we showed that apical SMA-1/βH-spectrin helps organize two specific AFBs within a set of four parallel AFBs present in the *C. elegans* L4 seam epidermis. In *sma-1* mutants, reduced and less coherent medial AFBs lead to altered seam mechanics, expanded regions of matrix delamination, reduced matrix adhesion, and partial or complete loss of the middle alae ridge (Fig. 9A). Thus, through its effects on the actin cytoskeleton, spectrin can also profoundly change the extracellular matrix and its patterns on animal surfaces.

**Figure 9.**
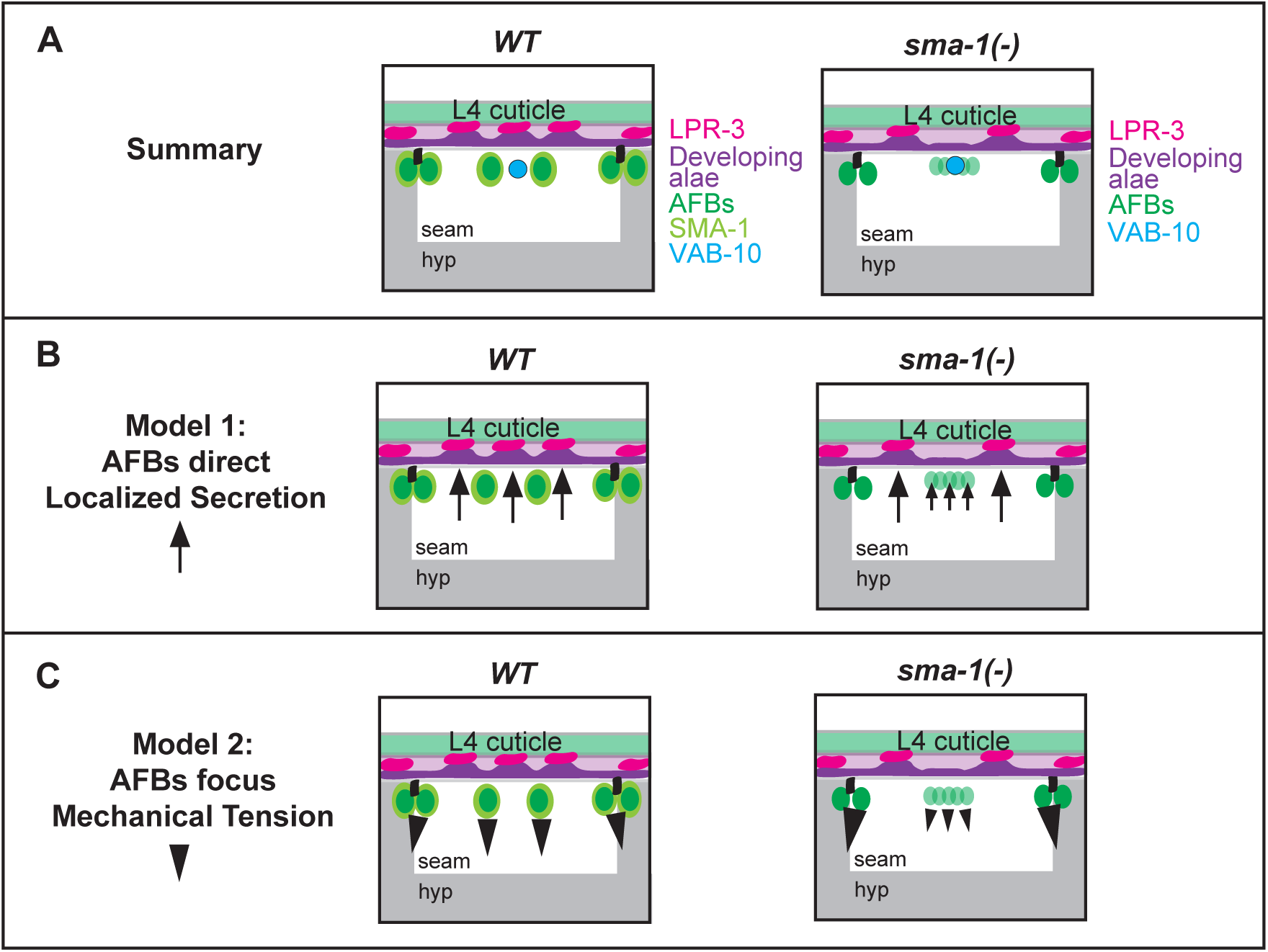
Models for SMA-1- and AFB-dependent patterning of alae ridges. (A) Results summary. Loss of *sma-1* reduces medial AFBs and leads to regions of middle ridge loss, without disrupting medial VAB-10. Note that, for simplicity, VAB-10 is not shown in subsequent panels. (B) Model 1: SMA-1-dependent AFBs direct localized trafficking, which becomes less focused in mutants. (C) Model 2: SMA-1-dependent AFBs focus mechanical tension, which becomes broader in mutants. See text for details.

### SMA-1/βH-dependent organization of the seam cortical cytoskeleton

*C. elegans* alae form above the lateral seam cells, which are narrow strips of syncytial epidermis that are less than five microns wide but close to a millimeter long. Seam syncytia sit nestled on top of the larger hyp7 syncytium, which surrounds them on three sides and to which they are mechanically coupled through adherens junctions (Fig. 1A,B). The seam’s elongated topology is quite different from that of most other epithelial cell types in which apical spectrin function has been studied. Seam syncytia also have an unusual cytoskeletal architecture, with four parallel longitudinal AFBs that are closely associated with SMA-1/βH-spectrin and a single central band of intermediate filaments and VAB-10/spectraplakin.

Within this unusual seam cytoskeleton, only the two medial AFBs require SMA-1 for their coherence. These medial AFBs lack the stabilizing influence of adherens junctions and of the basolateral spectrin network (UNC-70 + SPC-1) and therefore appear particularly reliant on interactions with the apical spectrin network (SMA-1 + SPC-1). Loss of *sma-1* reduces medial AFBs and specifically affects just the middle alae ridge, supporting local patterning of that ridge by the AFBs that immediately flank it. The observed expansion of the matrix delamination zone in these mutants indicates that SMA-1 and robust medial AFBs are not required for delamination, but rather focus such delamination to the proper regions (Fig. 9).

We originally hypothesized that SMA-1 might link medial AFBs to the apical membrane and thereby directly relay tension for delamination, but our results suggest this may not be a critical aspect of SMA-1 function. Structure/function studies showed that the SMA-1 membrane-binding PH domain is dispensable for its function in alae patterning, as are the SH3 domain and many of the spectrin repeats; only mutations that remove the actin binding domain have a strong effect on alae. Furthermore, medial regions of delamination are not only present in *sma-1* null mutants, but they are actually enlarged, indicating that SMA-1 restrains rather than promotes delamination. *sma-1* mutants also show increased TES-1 recruitment to junctional AFBs, suggesting that through its effects on actin, SMA-1 does influence seam mechanics even if it does not play a direct role in AFB-membrane attachment.

Seam medial AFBs are reduced but not entirely absent in *sma-1* mutants, and the medial VAB-10 network is still intact (Fig. 9A), suggesting some spatial information still remains, which may explain why a partial middle alae ridge is still present. Among the other factors contributing to seam cytoskeletal organization could be the ezrin/radixin/moesin homolog ERM-1 [31], another membrane-actin linker that our initial screen identified as having AFB-like localization patterns (Table S1). Further studies will be needed to understand the roles of ERM-1 and other actin-binding proteins present within the seam and their relationships to SMA-1.

### The role of VAB-10/spectraplakin in alae patterning

VAB-10/spectraplakin is another actin binding protein that influences alae development but whose role remains to be clarified. VAB-10 and its known partner IFB-1b have an intriguing localization pattern in the inter-AFB region below the central alae ridge, yet it’s not clear what role, if any, this pool of VAB-10 plays in alae patterning. This work shows that VAB-10a does not rely on SMA-1 for its medial recruitment, nor does it appear to act redundantly with SMA-1 to pattern the middle ridge, although seam-specific knockouts will be needed to fully address its role. Instead, the fact that *vab-10a(rf); spc-1(rf)* double mutants have defects in all three ridges suggests that both play larger, SMA-1-independent roles in actin organization or other processes. VAB-10 may be acting not only in the seam but also or instead in hyp7, which is mechanically coupled to the seam and has both longitudinal and circumferentially-arranged actin, spectrin, and VAB-10 structures. The contributions of the hyp7 cytoskeleton to alae patterning, already suggested in [11], will be important to dissect in future studies.

### How is seam AFB information relayed to the aECM to pattern alae?

While prior studies had suggested that seam AFBs pattern alae spacing and orientation [11], the extreme pleiotropy associated with knocking down actin or NM II made it challenging to directly link specific cytoskeletal changes to their matrix consequences. The specific consequences of removing SMA-1/βH-spectrin now reveal that the middle alae ridge is patterned independently from the outer ridges, by the flanking actomyosin structures below it.Furthermore, SMA-1-dependent medial AFBs restrain rather than promote delamination.

The mechanism by which AFBs influence delamination and other matrix properties remains an open question. One possible model is that AFBs direct localized secretion, such as by constraining matrix secretion to the regions between them to build the ridges (Fig. 9B, Model 1). In *sma-1* mutants, reduced medial AFBs could lead to less focused secretory activity and therefore less efficient adhesion and broader delamination (Fig. 9B). While such a patterned secretion-based model can’t be excluded, we have not found evidence to support such a model. Our ultrastructural data have not revealed periodic patterns in vesicle position or abundance. All pre-cuticle fusion proteins imaged to date (including LPR-3, which marks nascent ridges that correspond to regions of adhesion - see Fig. 1) appear broadly secreted prior to pattern emergence [11].

We previously proposed an alternative model that AFB information could be transmitted mechanically to the matrix via transmembrane matrix factors [11] (Fig. 9C, Model 2). Both *in vitro* studies and *in silico* modeling have shown that mechanical forces can pattern and deform fibrous collagen matrices over distances greater than several cell diameters [32–35]. When combined with the known tendencies of layered biomaterials to delaminate and wrinkle under mismatched strains and adhesiveness, AFB-dependent cues could lead to reproducible patterns of delamination vs. adhesion and matrix buckling. While our new results indicate that medial AFBs are not required for delamination, they do suggest that medial AFBs help to distribute forces within the tissue. In *sma-1* mutants, reduced medial AFBs and increased junctional stress may lead to altered strain on the matrix that favors delamination over adhesion.

Distinguishing between these (and other) models will require more direct measurements of seam mechanical properties and the identification of additional molecular players, such as other aECM factors that might be locally secreted in the case of Model 1, or other proteins that link AFBs to the membrane and/or the membrane to the matrix in the case of Model 2.Identifying such factors is a major goal of ongoing work.

### Actin binding proteins and the evolution of invertebrate surface structures

Invertebrates exhibit a wide variety of cuticle substructures that are thought to confer advantages in different ecological niches. For example, adults of different nematode species have different numbers or shapes of alae ridges or can lack alae entirely [36,37]. The functional consequences of alae differences are still a mystery, but the very specific *sma-1* mutant phenotype described here echoes classic studies on actin in Drosophila bristle cells [38,39] and reveals how changes in a single actin binding protein can lead to specific changes in cuticle patterning. We predict that such changes over the course of nematode evolution could explain many of the observed species-specific morphologies.

## Methods

### Strains and Worm Maintenance

C. *elegans* strains used in this study are listed in Tables S1 (fusions screened) and S3 (strains studied further). Fusion proteins were expressed from their endogenous loci, except for actin which was visualized using seam-specific transgenic reporters expressing the utrophin CH domain (UTRNCH) fused to GFP or dsRed [11]. Information from Wormbase was used in the design of all experiments [40]. Strains were maintained under standard conditions at 20°C on nematode growth medium (NGM) plates seeded with *E. coli* OP50-1 [41].

### Light Microscopy

Epifluorescent and differential interference contrast (DIC) images were collected with a Zeiss Axioskop (Carl Zeiss Microscopy). Confocal images were collected with a Leica Dmi8 microscope, HC PL APO 63x (Numerical Aperture 1.3), and Las X software. Super-resolution (Airyscan) images were collected with a Zeiss LSM 880 microscope, Airyscan Plan-Apochromat 63x/1 or Plan/LD C-Apochromat 40x/1.3/1.1 objective, and Zen Black software. In all cases, 400 Hz scanning speed, 1X line accumulation, 4X line average, and a pinhole setting of 1 AU were used. For confocal, Z stacks were collected with 0.33 μm slices and for Airyscan Z-stacks were collected in a range of 0.19-0.33 um slices. SfGFP fusions were excited via a 488 nm laser and emissions between 493 and 547 nm were detected using a HyD sensor. mCherry fusion proteins were excited by a 552nm laser and emissions between 583 and 784 nm were detected by a HyD sensor. Laser power (0.5-4%) and gain settings (60-90%) were determined based on fusion protein. Unless otherwise indicated, all confocal and Airyscan images shown are single Z-slices.

Live worms were immobilized with 10mM levamisole in M9 and mounted on a 5% agarose pad; for confocal and Airyscan imaging, pads were supplemented with 20mM sodium azide. Imaging of cytoskeletal protein fusions focused on the mid-L4 stages when alae patterning begins [11]. DIC imaging of alae was done in L4.9 larvae or young (non-gravid) adults. The precise stages of L4 worms were determined based on vulva morphology [42,43].

### Transmission electron microscopy

The TEM images of *sma-1(e30)* (CB30) adults in Fig. 3C are archival images from a prior study [44]. For TEM in Fig. 7, wild-type (N2) and *sma-1(ru18)* (AZ30) L4 worms were processed by high pressure freezing starting with 2%OsO4, 0.1%UAc, 2%H2O in acetone [45]. After freeze substitution, specimens were rinsed 4 times in 100% acetone, infiltrated into 2:1 and 1::2 acetone/Spurr resin, and then 4X in 100% Spurr resin prior to curing in a 60-degree oven for 2 days. Seven *sma-1(ru18)* L4 animals and five additional N2 L4 worms were cut at 70nm on an MRC PowerTome XL microtome. Sections were either post-stained in 2% Uranyl Acetate in H2O for 20min, followed by Reynolds lead staining for 3min, or not stained at all due to good membrane contrast. At least two thin sections from different mid-body regions per animal, separated by 20um along the body axis, were observed by electron microscopy. Either a TALOS L120C TEM (Thermofisher, Waltham MA) or a JEOL1400 Flash TEM (Peabody MA)with an AMT digital camera (Waltham MA.) were used. Images were processed in ImageJ [46] and manually pseudocolored in Adobe Illustrator (Adobe, San Jose California). We imaged a total of at least 10 specimens per genotype in order to find 3 N2 and 4 AZ30 mid-L4 specimens at the delamination stage. The N2 specimens were described in our prior study [11].

### DiI staining of cuticles

DiI staining to visualize alae was performed as previously described (Schultz and Gumienny 2012). Briefly, young adult worms were washed and pelleted from plates with M9 buffer and incubated in 30 µg/mL of lipophilic DiI stain (Invitrogen, Lumipore) shaking in the dark at 20 °C for 3 h. After a final M9 wash, the worms were allowed to recover on NGM seeded with OP50 in the dark for 30 minutes before mounting and imaging on the Zeiss Axioskop.

### Image Analysis

Alae phenotypes in Figs. 3-5 were scored by DIC imaging of late L4 larvae or young (pre-gravid) adults using the rubric described in the main text. Image measurements and quantifications were performed with ImageJ [46]. Fluorescence intensity quantifications in Fig. 6 and Fig. 8 used the box Measure tool to calculate total fluorescence within a 1X1 micron square box in the medial seam and/or the Plot Profile tool to measure peak fluorescence at junctional AFBs. Seam width (Fig. S3) quantifications used the line Measure tool. For TEM images, matrix slit width quantifications (Fig. 7D) used the line Measure tool. Where possible, measurements were made of the four slits on both the left and right sides of two different body regions per specimen (yielding up to 16 measurements per specimen); however, because the precise timing of slit formation varies along the body, some sides or regions did not yet have slits present to measure.

Statistical analysis was carried out using GraphPad Prism. For categorical data presented in Fig. 3F, 4B and 5C, phenotypic categories were combined so that outcomes were classified as normal vs abnormal or severe vs other categories for the Fisher’s Exact Test.

### RNA-mediated interference (RNAi)

Bacterial-mediated feeding RNAi was adapted from [47]. To knock down *vab-10, sma-1,*and *act-2*, we used sequence-verified plasmids from the Ahringer library. Single colonies of the *E. coli* strain HT115 (DE3) containing these plasmids were grown in LB and induced with isopropyl β-D-1-thiogalactopyranoside (IPTG) after ∼5-6 hours (1mM for *vab-10*; 3mM for *sma-1;* 8mM for *act-2*) and incubated for a further 1-2 hours before seeding on NGM plates supplemented with 1mM Amp and 1mM (*vab-10*), 3mM (*sma-1*), or 8mM (*act-2*) IPTG. As a control, we used *E. coli* HT115 (DE3) transformed with the empty vector L4440/ pPD129.36 that produces short dsRNA molecules that do not match any annotated gene of *C. elegans.* Single L4 hermaphrodites were placed per plate and transferred to fresh plates after 24 hours. The day 2 progeny were imaged by DIC as young adults (*vab-10*, *sma-1)* or by confocal microscopy as L4s (*act-2*). Both *vab-10* and *act-2* RNAi caused >50% embryonic or early larval lethality under our conditions, so only escapers could be imaged. When present, animals with a small size were prioritized for imaging since they were clearly affected by the RNAi.

Tissue-specific RNAi used the *rde-1(ne300)* null mutant background [48] with tissue-specific *rde-1+* rescue transgenes *aaaIs4 (elt-5pro::rde-1+)* [11] or *mfIs70 (lin-31pro::rde-1+)*[26] (see Table S3 for strains).

### Data and Resource Availability

All relevant data are contained within the figures, tables, and supplemental materials. Strains are available at the *Caenorhabditis* Genetics Center (U. Minnesota) or by request from the corresponding author.

## Acknowledgements

We thank Nathalie Pujol, Erfei Bi, Paul Janmey, Vivek Shenoy, and members of our lab and of the UPenn worm group for helpful discussions and advice; Matthew Buechner for sharing archival CB30 TEM images; Andrea Stout and the UPenn CDB Microscopy Core (RRID SCR_022373) for training and help with confocal and super-resolution microscopy; Biao Zhao and the UPenn TEM core for training and help with electron microscopy; and Jeff Hardin, Bob Goldstein, and Michel Labouesse for strains. Some strains were obtained from the *Caenorhabditis* Genetics Center (U. Minnesota), which is funded by the NIH Office of Research Infrastructure Programs (P40 OD010440). This work was funded by NIH grants R35 GM136315 to M.V.S., OD010943 to D.H.H., and by funds from the Portuguese Foundation for Science and Technology (FCT) under the project UIDB/04293/2020 as well as FCT funds to A.X.C (CEECIND/01967/2017 and 2023.14140.TENURE.013) and to F.Y.C. (DL 57/2016/CP1355/CT0013).

## Notes

### Competing Interest Statement

The authors have declared no competing interest.

